# Dielectrophoretic separation of randomly shaped protein particles

**DOI:** 10.1101/2020.07.23.218438

**Authors:** Tae Joon Kwak, Huihun Jung, Benjamin D Allen, Melik C Demirel, Woo-Jin Chang

**Affiliations:** Nancy E. and Peter C. Meinig School of Biomedical Engineering, Cornell University, Ithaca, NY, USA; Center for Research on Advanced Fiber Technologies, Materials Research Institute, Pennsylvania State University, University Park, PA, USA; Department of Engineering Science and Mechanics, Pennsylvania State University, State College, PA, USA; Department of Biochemistry and Molecular Biology, Pennsylvania State University, University Park, PA, USA; Huck Institutes of the Life Sciences, Pennsylvania State University, University Park, PA, USA; Department of Mechanical Engineering, University of Wisconsin-Milwaukee, Milwaukee, WI, USA; School of Freshwater Sciences, University of Wisconsin-Milwaukee, Milwaukee, WI, USA

**Author notes:** Corresponding authors. M.C.D.: 212 Earth and Engineering Science Bldg, Pennsylvania State University, University Park, PA, 16802, USA. W.-J.C.:3200 N Creamer St, EMS1113, University of Wisconsin-Milwaukee, Milwaukee, WI, 53211. This study was mostly done while T.J.K. was working at University of Wisconsin-Milwaukee. T.J.K. and H.J. contributed equally to this work.

**Keywords:** Protein particle separation, Morphology-based separation, Size-based separation, Dielectrophoresis, Protein particles, Protein aggregates

## Abstract

Recently, insoluble protein particles have been increasingly investigated for artificial drug delivery systems due to their favorable properties, including programmability for active drug targeting of diseases as well as their biocompatibility and biodegradability after administration. One of the biggest challenges is selectively collecting monodisperse particles in desirable morphologies and sizes to enable consistent levels and rates of drug loading and release. Therefore, technology that allows sorting of protein particles with respect to size and morphology will enhance the design and production of next-generation drug delivery materials. Here, we introduce a dielectrophoretic (DEP) separation technique to selectively isolate spherical protein particles from a mixture of randomly shaped particles. We tested this approach by applying it to a mixture of precipitated squid ring teeth inspired tandem repeat protein particles with diverse sizes and morphologies. The DEP trapping system enabled us to isolate specific-sized, spherical protein particles out of this mixture: after separation, the fraction of 2 μm and 4 μm spherical particles was increased from 28.64% of mixture to 80.53% and 74.02% with polydispersity indexes (PDIs) decreased from 0.93 of mixture to 0.19 and 0.09, respectively. The protein particles show high aqueous swelling capability (up to 74% by mass) that could enable delivery of drug solutions. This work is intended to inspire the future development of biocompatible drug-delivery systems.

## 1. Introduction

Bio-derived materials are an enduring feature of human experience. Many bio-derived materials, such as silk, collagen, and wool, are made of proteins with repetitive amino acid sequences [1]. Tandem repeat sequence segments, or motifs, play key roles as the building blocks of structures and functions in these protein materials [2, 3]. Tandem repeat protein evolved via gene duplication, which is a major mechanism for new structures and functions. The potential for programmable biosynthetic materials has attracted increasing attention due in part to the discovery of these motifs. By combining the concept of biomimicry and theoretical foundations of block copolymers, these structural motifs have been exploited to design protein-based materials with remarkable new properties. Many reports have shown that these new materials can have interesting and useful mechanical, optical, and thermal properties [4–11]. Moreover, tandem repeat proteins govern the assembly of composites materials such as graphene oxide and MXenes (Ti_3_C_2_T_x_) for 2D layered structures as well as self-healing characteristics of conducting polymers for applications in flexible electronics and biocompatible electronics [12–14].

Given the recent development of programmable protein materials, several applications to artificial drug-delivery systems have been reported [15, 16]. Crucial requirements for carrier materials in drug-delivery applications include high and constant loading of therapeutic agents, maintainance of physical structures and physicochemical properties [17] in the patient body during drug delivery, and harmless degradation after the delivery process is finished. To meet these requirements, a variety of separation methods have been used to isolate protein particles that have the desired properties. Centrifugation has been used with the sucrose-gradient technique to separate produced protein particles from non-compacted particles by sedimentation velocity [18]. Gel filtration chromatography is used to separate proteins by size and under native or denaturing conditions using porous gels [19]. In addition, various chromatography methods that utilize molecular interaction, affinity, and ion exchange are widely being used [20]. However, these approaches still remain challenging in the separation of proteins with the consistency of both size and morphology from all others.

Dielectrophoresis (DEP), an electrical particle-manipulation technique, has been utilized for a variety of biomedical applications such as particle separation and patterning [21–23], intermolecular force spectroscopy [24, 25], localized heat generation [26], and drug delivery and discovery [27–29]. When dielectric particles are introduced into a non-uniform electric field, these particles experience electrical attraction or repulsive force according to the physical/electrical properties of the particles and media, and distribution of the electric field gradient, which is called the DEP force. Notably, DEP has been widely used for the manipulation of particles in various biological and biomedical applications because it offers advantages such as easy operation, high-throughput efficiency, tighter control, and less damage to the particles, compared with traditional techniques such as filtration, centrifugation and electrophoresis [30, 31].

Here, we demonstrate dielectrophoretic separation of designer protein particles based on size and morphology within a microfluidic DEP trapping device in a single operation. The capability of the protein particles to deliver aqueous solvents was tested using a mixture of water and urea as a model. The protein powder was produced using recombinant expression in *E. coli* bacteria followed by single step purification. From the powder, protein aggregates with a distribution of sizes and morphologies were synthesized by precipitation from a solution of hexafluoroisopropanol, one of the few solvents that can dissolve this type of protein. To facilitate the separation of this diverse mixture, we fabricated an array of circular openings on a chromium layered glass substrate with an indium-tin-oxide (ITO) counter electrode to apply an omni-directional DEP force to the center of a circular trap. The system allows the selective isolation of specific size of spherical protein particles from the diverse mixture.

## 2. Materials and Methods

### 2.1. Gene editing, protein expression and extraction

Construction of repetitive DNA constructs, their expression in *E. coli* and purification were well described in our previous work [6]. Briefly, unit gene (n=1) fragments including cloning sequences were purchased from Genewiz (Figure 1a). Double-stranded templates for protected digestion of rolling circle amplification (PD-RCA) were prepared by ScaI digestion, followed by blunt-end ligation. PD-RCA reaction was performed for 24 hours (18 hours of RCA and 6 hours of PD), and then agarose-gel-purified products (Omega Bio-Tek E.Z.N.A gel extraction kit) were cloned into the open-reading frame of an expression vector (pET14b). Sequence-verified plasmids (by Sanger-sequencing at Genomic core facility at Pennsylvania State University) were transformed into the BL21 (DE3) expression strain.

**Figure 1.**
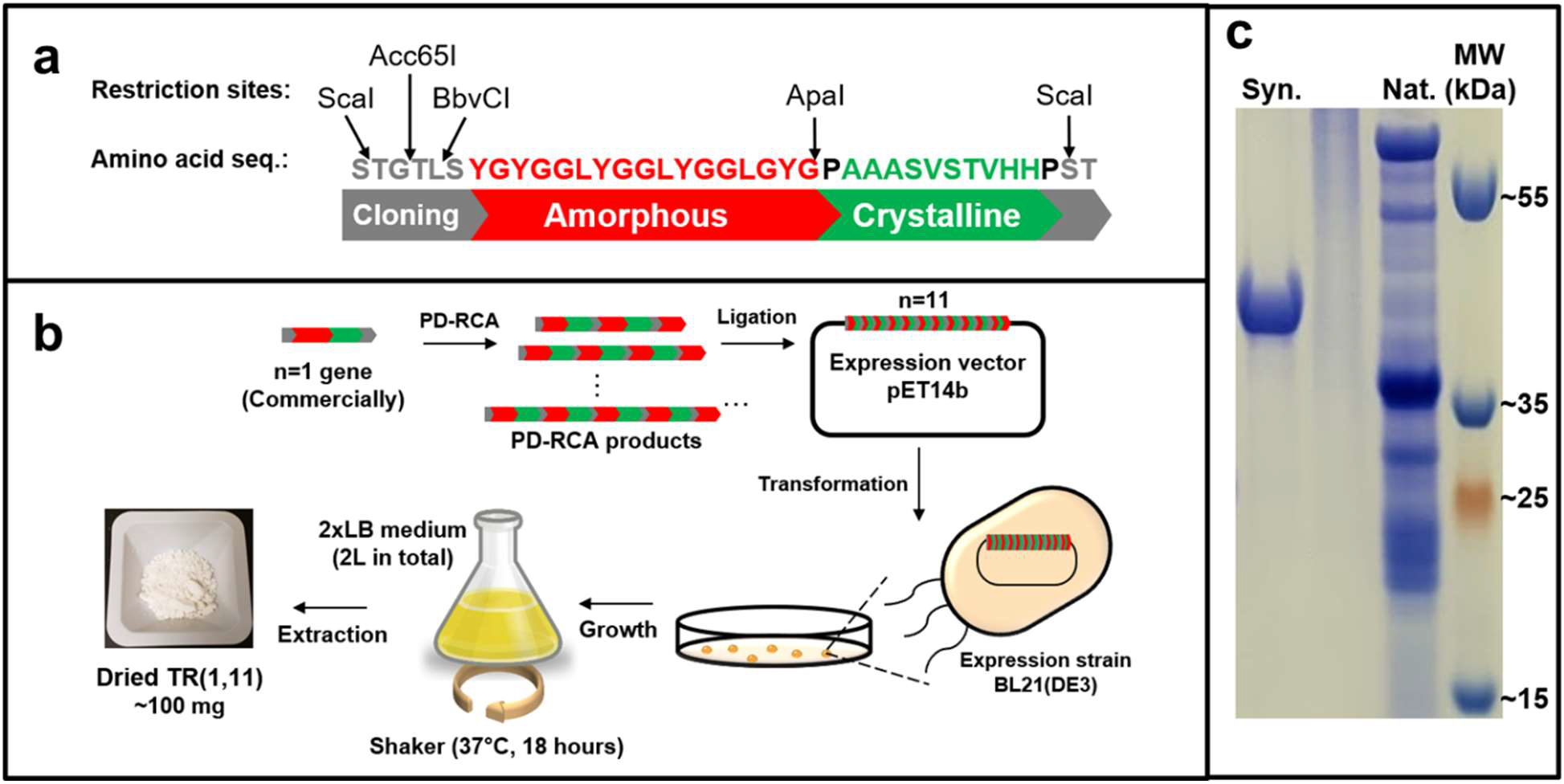
Genetic information, schematics and molecular information of the protein material. (a) the unit sequence of the protein with cloning sites for repetitive modification, (b) schematic illustration of entire process from gene construction to protein production, and (c) an SDS-page gel image showing the difference in molecular weight between the designed protein material and native squid ring teeth proteins (SRTs). The unit sequence of the designed protein was inspired by the native SRTs in this gel image.

2x Luria-Bertani (2xLB) broth was chosen as bacterial culture media (20 g of tryptone, 10 g of yeast extract, and 10 g of sodium chloride per liter of double distilled water). A single colony was used to inoculate 100 mL of LB-Miller incubating at 29°C and 210 rpm for about 10 hours as a starter culture (OD600 = 0.5 – 1, pH = ~7). After this incubation, the starter culture was added to 2 L of 2xLB liquid medium and grown at 37°C and 300 rpm until OD600 reached around 3 (pH = ~7). The 2-L culture was induced with isopropyl β-D-1-thiogalactopyranoside (IPTG) of 1 mM as a final concentration for 8 hours, and then harvested by centrifugation (10,000 rpm for 15 minutes, 12 - 15 gram of wet mass in total). For protein extraction, re-suspended cell pellets (50 mM Tris pH 7.4, 200 mM NaCl, and 2mM EDTA, 3 gram of wet mass pellet per 250 mL) were lysed by high-pressure homogenizer (Microfluidizer, Microfluidics-M110EH-30) at 15,000 psi, and lysed pellets were washed with urea-extraction buffer (100 mM Tris pH 7.4, 5 mM EDTA, 2 M Urea, 2% (v/v) Triton X-100, 100 mL per 3 gram of wet pellet lysate) and last-extraction buffer (100 mM Tris pH 7.4, 5 mM EDTA, 100 mL per 3 gram of wet pellet lysate, twice wash) to remove left-over cell debris generated during physical extraction process by low-spin centrifugation (5,000 rpm). Washed pellets were lyophilized with FreeZone 6 plus (Labconco) for 12 hours at CSL Behring fermentation facility at Pennsylvania State University. The yield of protein expression was approximately 200 mg per liter of 2xLB liquid medium.

### 2.2 Protein particle processing

Lyophilized purified protein was completely dissolved into hexafluoroisopropanol (HFIP) to a final concentration of 20 mg/mL. The protein-HFIP solution was slowly diluted 10-fold into 50mM sodium dodecylbenzenesulfonate (NaDBS), and mixed thoroughly by pipetting several times. Proteins began to precipitate immediately. To stabilize the protein particles, the suspension of precipitated protein was stored at room temperature for several hours. The protein particles were washed with double-distilled water (ddH_2_O) twice by low speed centrifugation (at 8000 rpm for 3 minutes) to remove HFIP and NaDBS. The final conductivity of the protein particles in ddH_2_O was 2 μS·cm^−1^.

### 2.3. Microfluidic trapping device fabrication and experimental setup

The microfluidic trapping device for the DEP separation was fabricated using conventional photolithography and metal-deposition processes. Figure 2 illustrates the experimental setup and concept of the lab-on-a-chip system. A circular trap array pattern was prepared on a 1-inch by 1-inch glass microscope slide through the lift-off process, where each circular trap was 20 μm in diameter with a 10-μm gap between the two adjacent circular traps. A 100-nm thick chromium layer was deposited using a thermal evaporator (KV-301, Key High Vacuum Products, Inc) and patterned by a lift-off process. A 30-μm-high microfluidic flow channel was made using a polyethylene terephthalate (PET) spacer (Nitto Denko Co) and an ITO-coated glass slide. Flow inlet and outlet holes on the glass substrate were pierced by a highspeed rotary tool with a diamond-coated drilling tip. Through these holes, the liquid media carrying the particles was injected at the inlet using a syringe pump (KDS 200, KD scientific Inc), and drained at the outlet via a capillary tubing. The chromium DEP electrode and ITO counter electrode were connected to an arbitrary waveform generator (33250A, Agilent) and the signal was measured using a digital storage oscilloscope (2190D, B+K Precision) in real-time. Real-time observation and image captures were accomplished using a cooled charge-coupled device camera (DP-72, Olympus) equipped on an upright microscope (BX53F, Olympus).

**Figure 2.**
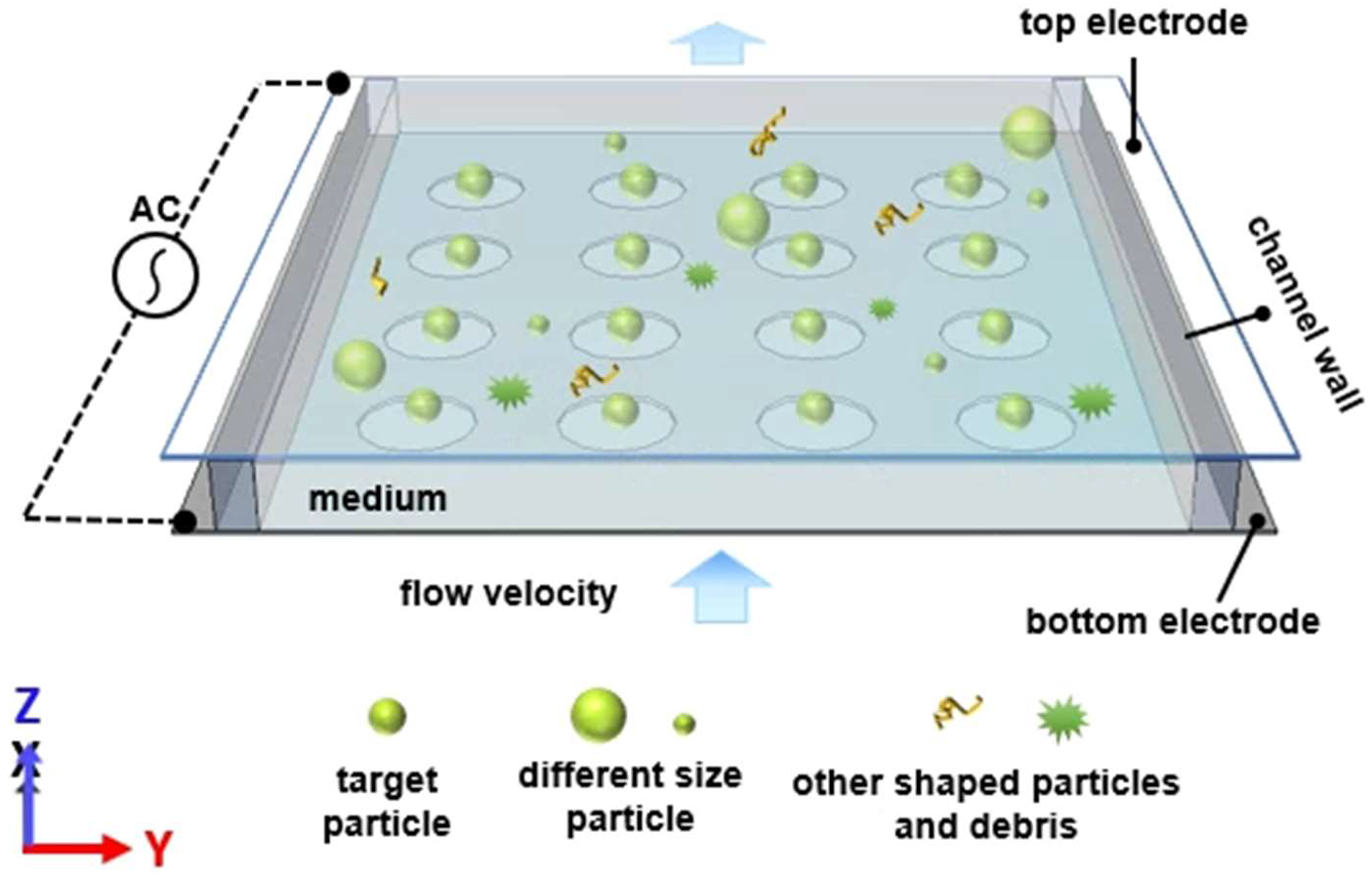
Schematic illustration of the lab-on-a-chip system and experimental setup. In the lab-on-a-chip system, target particles are separated from the mixture of other particles depending on size and morphology.

### 2.4. Numerical analysis

To determine optimum condition for particle trapping, the characteristics of the circular traps were analyzed using finite element analysis (FEA) with numerical simulations using Comsol Multiphysics® (v5.2, COMSOL Inc) with the AC/DC module. By the numerical analysis, the geometrical effect of the trap electrode, electric field distribution, electrical potential, and magnitude and direction of the DEP force were investigated. The size of the mesh was initially set to be less than 200 nm around and inside the trap, and was interpolated at intervals of 10 nm around the particle in order to obtain a uniform interval resolution in the numerical analysis according to the particle positions. The DEP force acting on an individual particle was calculated by using an equation:

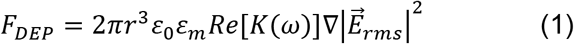

where *ε*_0_ and *ε*_*m*_ are the permittivity of the free space and medium, respectively, *E* is the applied electric field, and *K*(*ω*) denotes the Clausius-Mossotti (CM) factor which contains all the frequency dependence and the components of the complex dielectric constant of the DEP force. The CM factor is expressed as:

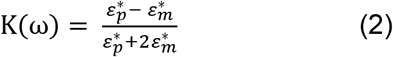

here, the indices *p* and *m* denote the permittivity of the particle and the surrounding medium, respectively.

### 2.5. Fourier transform infrared spectroscopy (FTIR)

Experiments were performed with a Shimadzu IRTracer-100 under attenuated total reflection (diamond crystal) mode with 2 cm^−1^ resolution from wavenumber 450 to 4000 cm^−1^. 256 scans were co-added for each measurement to collect enhanced-resolution spectra. For the secondary-structural-content calculation, deconvolution of the spectra was performed using OriginPro 8.5 software. The amide I band (1580 – 1720 cm^−1^) was selected for deconvolution after linear baseline correction of both end points. The peak position of each deconvoluted curve was determined by the local minima of second-derivative spectra [32], followed by fitting using a nonlinear least-squares method with a maximum of 500 iterations. Each deconvoluted curve was assigned to secondary-structural components [33–43], and their percentages were calculated by summation of total area under assigned curves.

### 2.6. Thermogravimetric analysis (TGA)

TGA was performed on a TA Instruments Q5500 in order to measure hydration rates of spherical protein particles after captured by DEP chips. For analysis at full hydration, captured protein particles were immersed separately in three different solutions (water, 26.5% weight-volume urea solution, 39.8% weight-volume urea solution) overnight. While transferring fully-hydrated protein particles, one end of each wet protein sample was blotted with tissues for 5 minutes to remove excess solution on its surface. The initial sample weight was in the range between 5 mg and 15 mg.

## 3. Results and discussion

### 3.1. Protein particle characterization

To characterize the morphology and determine the size of the protein particles, the prepared stock dispersion (Figure 3a) was examined by scanning electron microscopy (SEM) and dynamic light scattering (DLS) measurement. As shown in Figure 3b, the morphology of the protein particles investigated by SEM (JEOL 6460LV, JEOL Ltd.) shows irregular shapes and dispersed sizes. We subsequently analyzed the size distribution of individual protein particles using DLS (NanoBrook 90Plus PALS, Brookhaven Instruments) measurement as shown in Figure 3c, and diameters from 1 μm to 26 μm were obtained. The average diameter was 2.91 μm with a PDI of 0.93, as the PDI value is close to 1, the individual protein particles have a heterogeneous size distribution. In addition, we analyzed the circularity of 439 protein particles using ImageJ image analysis software (v1.52a) as shown in Figure 3d. The average circularity index was 0.64 with a standard deviation of 0.21; the circularity index value far from 1 indicates that the shapes of the protein particles diverge widely from spherical.

**Figure 3.**
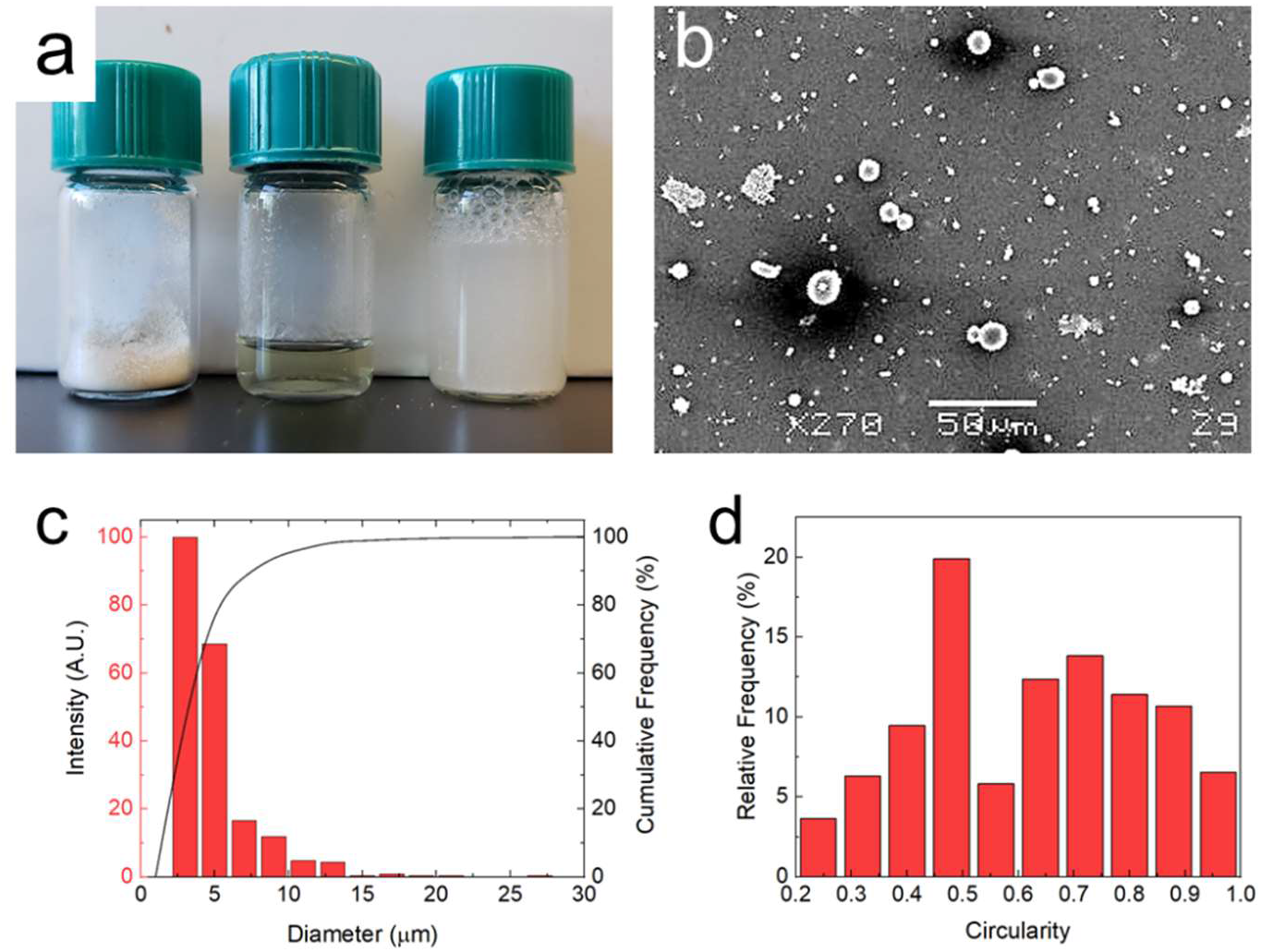
Particle characterization. (a) Picture of vials including powder (left), hfip solvent solubilized (center), final bead solution (right). (b) The morphology of the synthesized protein particles was examined by scanning electron microscopy imaging. Scale bar = 50 μm. (c) The size distribution of the synthesized protein particles by dynamic light scattering measurement. (d) The circularity distribution of the synthesized protein particles.

### 3.2. Separation of specific sizes spherical protein particles by DEP

Figure 4a presents numerically simulated direction and magnitude of the DEP forces around a nine-trap pattern array when voltage (i.e., AC 1 MHz 10 V_pp_) is applied to the electrodes. The simulation was performed using the geometry of the device, and experimentally determined permittivities and conductivities of the protein particles and media. Each arrow represents the negative DEP force vector at the location. The negative DEP force around the circular trap is radially symmetric toward the center of the trap, where the electric field gradient is minimum. Thus, the protein particles located inside of the trapping area will be pushed/pulled by the negative DEP force and become trapped.

**Figure 4.**
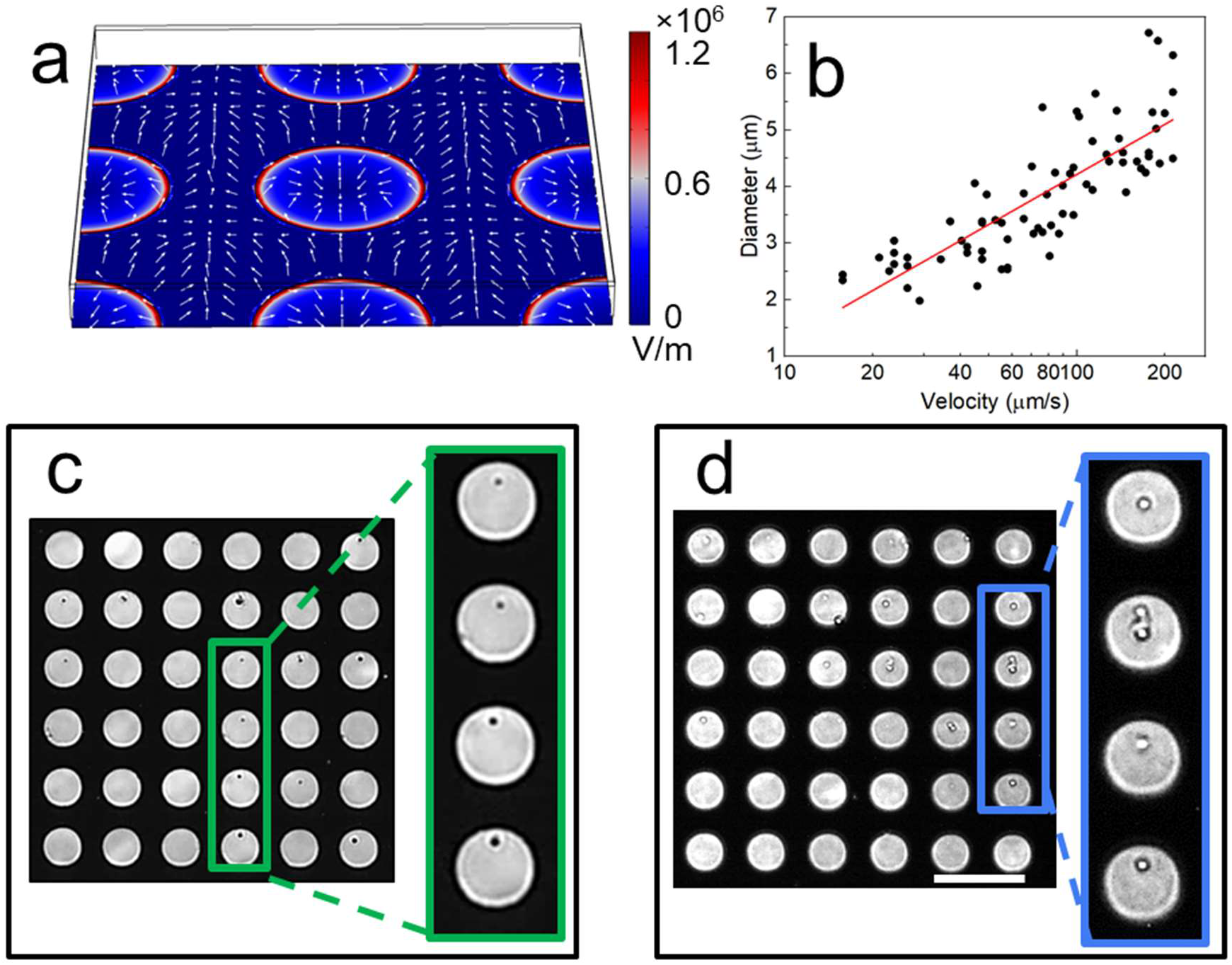
Dielectrophoretic separation of protein particles. (a) Numerical simulation by finite element method shows the magnitude distribution of the electrical potential (surface color) and direction of DEP force around the circular traps (arrows). Once the particle enters the circular trap, DEP force will pull and push the particle toward the center of the trap. (b) Velocity and size of the trapped protein particles when AC 1 MHz 10 V_pp_ was applied to the electrodes. (c and d) Captured images of the trapped ~2- and ~4-μm-diameter protein particles at two different flow rates of 25 μm/s and 70 μm/s, respectively. The enlarged images are displayed on the right side. Scale bar = 50 μm.

The protein particle trapping experiments were performed by introducing a stock solution of protein particles with diverse sizes and shapes suspended in ddH_2_O into the device; results are plotted in Figure 4b. The voltage applied to the electrodes was fixed at AC 1 MHz 10 V_pp_ during the experiment, and the flow velocity was controlled by the syringe pump. The velocity and size of the trapped protein particles are correlated as y = −1.66 + 2.937*×* (R^2^ = 0.66) with the logarithmic velocity axis. This result suggests that particles are trapped depending on size as well as velocity. Thereafter, two different flow rates of 25 μm/s and 70 μm/s were tested to trap ~2- and ~4-μm-diameter protein particles, respectively, with a fixed AC voltage of 1 MHz 10 V_pp_. Interestingly, only single-sized spherical-shaped particles were trapped from the mixture as shown in Figure 4c and d.

### 3.3. Characterization of protein beads after DEP separation

We subsequently performed morphological analyses on the ~2-μm and ~4-μm protein-particle samples collected by DEP separation in Figure 4c and d. The characterization conditions were the same as those used in Figure 3. The sizes of the separated protein particles are shown in Figure 5a and c. The average diameters of the trapped protein particles were 2.28 with a PDI of 0.19 and 3.79 μm with a PDI of 0.088, respectively. As the PDI values are close to 0, the trapped protein particles are monomodally dispersed with respect to size (Figure 5a). Moreover, the circularities of the trapped particles are shown in Figure 5b and d, and the average circularity indexes were 0.87 ± 0.1, and 0.81 ± 0.19, respectively. As the circularity index value is closer to 1, the protein particles conform to a spherical shape. This means the DEP trapping system can selectively capture spherical particles and exclude non-spherical particles. This can be explained by the effect of the folding factor, where the effect of the DEP force acting on the particle increases when the shape of the particle is closer to a sphere [44, 45]. Although the dimensions of the non-spherical protein particles are similar to the spherical ones, the reduced DEP forces on the non-spherical particles prevent trapping, and thus only the sphere-like protein particles are trapped. Our experimental results suggest that reduced DEP forces are applied to non-spherical particles regardless of whether the particles are larger or smaller (See supporting Video M1 and M2). Consequently, the sum of the relative frequencies from 0.8 to 1.0 in the circularity test for raw protein particle samples and the trapped ~2-μm and ~4-μm particles was increased from 28.64% of mixture to 80.53% and 74.02%, respectively.

**Figure 5.**
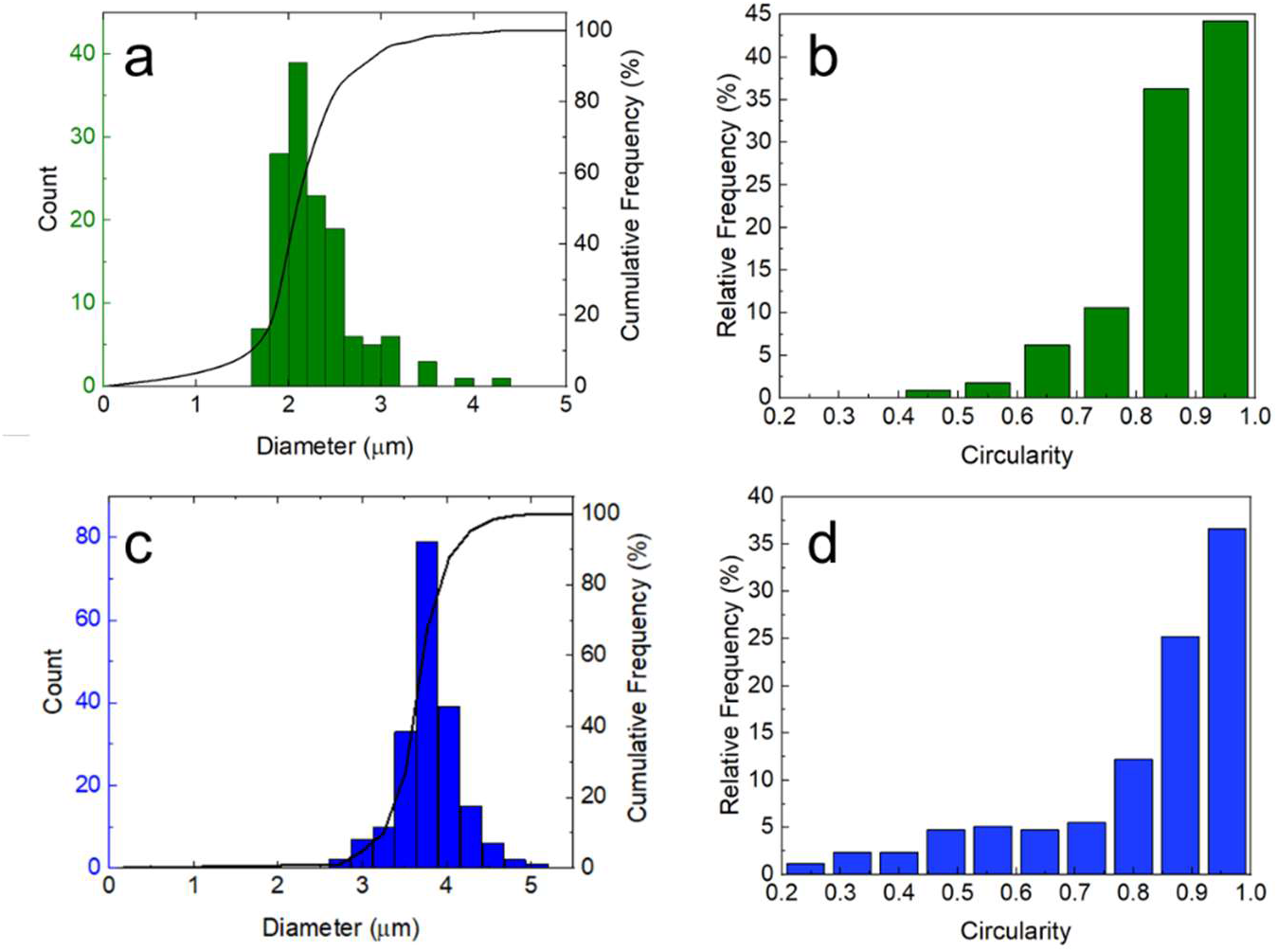
Morphological characterization of the trapped ~2-μm (a, b) and ~4-μm (c, d) protein particles. (a, c) The size distribution measurement of the trapped protein particles. (b, d) The circularity distribution of the trapped protein particles.

Secondary structure analysis was performed on spherical protein particles collected from DEP separation by Fourier-transform infrared spectroscopy (FTIR). Figure 6 clearly shows that insoluble particles separated by DEP are protein particles. Two strong Amide bands, Amide I (1600 – 1700 cm^−1^) and Amide II (1510 – 1580 cm^−1^) (Figure 6a) were detected. The peak-center position of the Amide I band for these samples occurs at 1530 cm^−1^, suggesting that β-sheet structures are dominant. By deconvolution of the Amide I band as shown in Figure 6b, quantitative analysis of secondary-structure percentage (Table 1) indicated that 58% of secondary structure in the captured protein particles are β-sheet, consistent with the prior results for these proteins [6].

**Figure 6.**
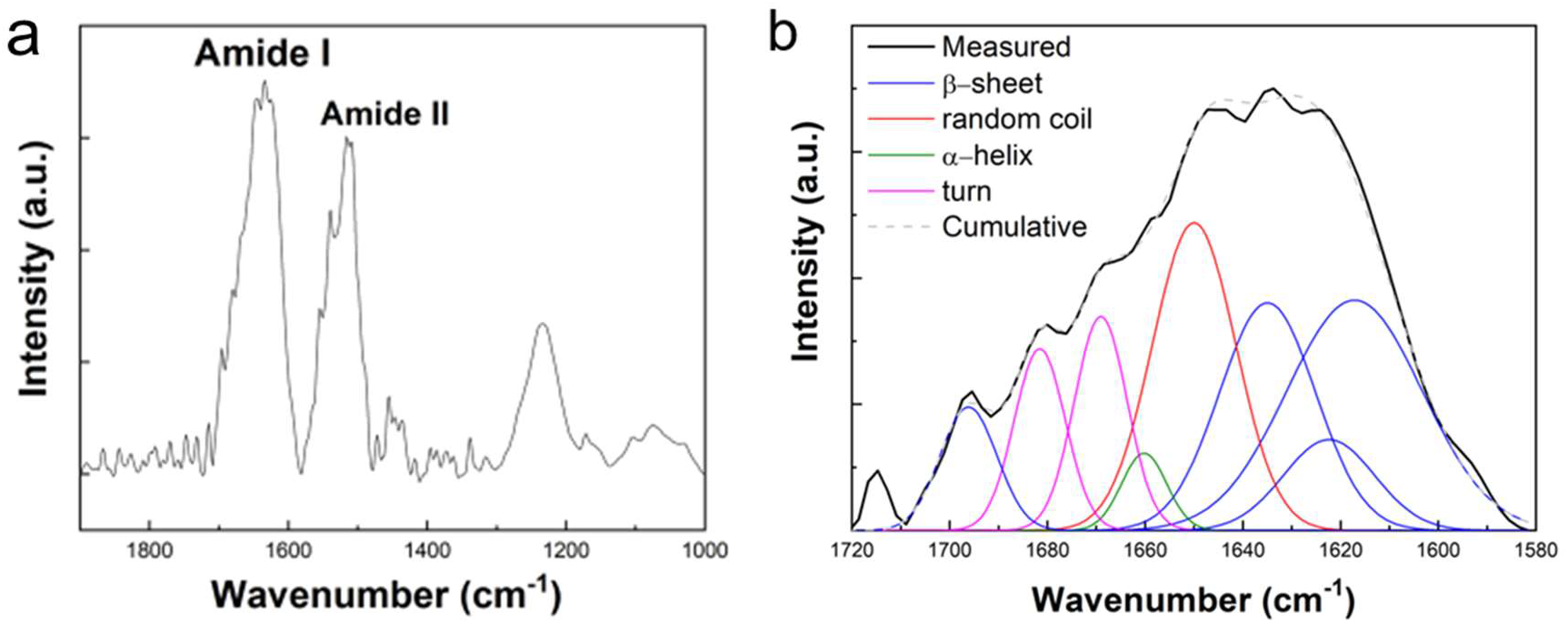
Fourier-transform infrared spectroscopic (FTIR) characterization of spherical-shaped protein particles trapped by DEP chips. (a) full spectrum, and (b) deconvolution of the Amide I band.

**Table 1.** Secondary-structure percentage of the spherical-shaped protein particles separated by DEP, as calculated by total portion of area under the fitted curves.

| Secondary structure | β-sheet | Random coil | Turns | α-helix |
| --- | --- | --- | --- | --- |
| **Percentage (%)** | 58 | 22 | 17 | 3 |

Hydration characteristics were tested for spherical protein particles separated by DEP by thermogravimetric analysis (TGA). Interestingly, TGA plots of protein saturated with urea solutions (Figure 7 red and blue curves) have two stages of mass loss near 110°C and 140°C. Water and urea evaporation occurred at 110°C and 140°C, respectively, due to the boiling point of water (100°C) and melting point of urea (135°C). TGA traces of mass loss for three different concentrations of aqueous solutions are shown in Table 2. Total hydration rate of the materials (Figure 7 and Table 2) was about 74% (72.96%, 73.59% and 75.28%, respectively, in three different solvent conditions). Calculations of urea concentration in solvents carried by protein particles are shown in Table 2. Urea concentrations carried by protein particles matched with those of original urea loading solvents. This clearly shows that protein particles can carry solutes.

**Figure 7.**
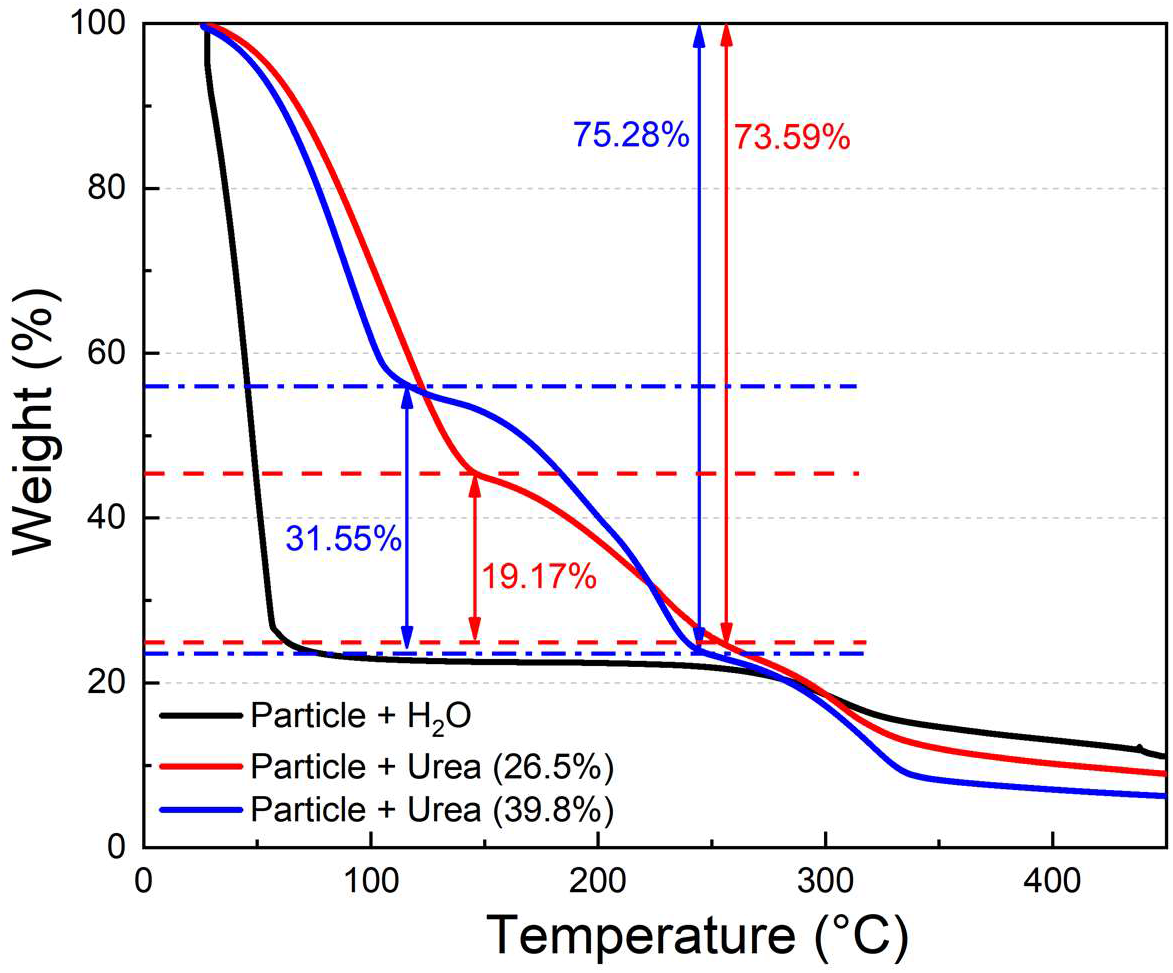
Hydration characteristics of separated protein particles. TGA curves with three different show carrying capacity of captured protein particles. The concentrations of solutes carried by protein materials were similar to those of the loading solutions.

**Table 2.** Percentage mass loss of protein particles with three different solutions

|  | Mass ratio<br>of water (%) | Mass ratio<br>of urea (%) | Mass ratio<br>of protein (%) | Urea portion (%) | Original urea<br>concentration<br>(%) |
| --- | --- | --- | --- | --- | --- |
| Water only,<br>black curve | 72.96 | 0 | 27.04 | 0 | 0 |
| Urea (26.5%),<br>red curve | 26.41 | 47.18 | 26.41 | $\frac{19.17}{73.59} = 26.05$ | 26.5 |
| Urea (39.8%),<br>blue curve | 31.55 | 43.73 | 24.72 | $\frac{31.55}{75.28} = 41.91$ | 39.8 |

## 4. Conclusion

In this study we produced protein particles and selectively collected spherical 2 and 4-μm particles from a mixture with diverse sizes and shapes using a DEP trapping system. In addition, we validated the capability of these size- and shape-separated protein particles to store and release a solute in aqueous solution. The results show the capability of rapid one-step separation of protein particles by DEP.

The high swelling ratio of the size- and shape-selected protein particles suggests that they may be able to deliver large amount of aqueous solutions. The ability to rapidly separate protein particles with desired sizes and shapes will enhance the rapid development and assessment of future protein-based drug carriers.

## Acknowledgments

We acknowledge the use of instrumentation at the Advanced Analysis Facility of the University of Wisconsin-Milwaukee. The authors also acknowledge Georgije Stanisic’s data analysis help. The authors are also grateful for the Distinguished Graduate Student Fellowship and Distinguished Dissertation Fellowship provided to T.J.K. by the University of Wisconsin-Milwaukee. MCD, HJ, BA are funded by the Army Research Office (grant no. W911NF-16-1-0019 and W911NF-18-1-0261) as well as Huck Endowment of Pennsylvania State University.

## Author contributions

T.J.K. and H.J. designed this research. T.J.K. and H.J. set up the instruments and performed experiments. T.J.K. carried out the numerical analysis with finite element simulation. T.J.K., H.J., M.C.D., and W.-J.C. analyzed the data. T.J.K., H.J., B.D.A., M.C.D., and W.-J.C. contributed to editing and valuable discussions on this manuscript, and all authors contributed to writing the manuscript.

